# M2 Macrophage Exosomes Reverse Cardiac Functional Decline in Mice with Diet-Induced Myocardial Infarction by Suppressing Type 1 Interferon Signaling in Myeloid Cells

**DOI:** 10.1101/2024.09.13.612924

**Authors:** Martin Ng, Alex S. Gao, Tuan Anh Phu, Ngan K. Vu, Robert L. Raffai

## Abstract

Effective treatment strategies to alleviate heart failure that develops as a consequence of myocardial infarction (MI) remain an unmet need in cardiovascular medicine. In this study, we uncovered that exosomes produced by human THP-1 macrophages cultured with the cytokine IL-4 (THP1-IL4-exo), reverse cardiac functional decline in mice that develop MI as a consequence of diet-induced occlusive coronary atherosclerosis. Therapeutic benefits of THP1-IL4-exo stem from their ability to reprogram circulating Ly-6C^hi^ monocytes into an M2-like phenotype and suppress Type 1 Interferon signaling in myeloid cells within the bone marrow, the circulation, and cardiac tissue. Collectively, these benefits suppress myelopoiesis, myeloid cell recruitment to cardiac tissue, and preserve populations of resident cardiac macrophages that together mitigate cardiac inflammation, adverse ventricular remodeling, and heart failure. Our findings introduce THP1-IL4-exo, one form of M2-macrophage exosomes, as novel therapeutics to preserve cardiac function subsequent to MI.

## Introduction

Heart failure (HF) remains a leading cause of morbidity and premature death worldwide, with a significant portion of these events resulting from hyperlipidemia-driven cardiovascular disease (CVD) (1). Occlusive coronary circulation, including via thrombosis due to atherosclerotic plaque rupture, results in cardiac ischemia and myocardial infarction (MI) (2). While survival rates from MI have improved owing to advances in interventional cardiology, long-term outcomes remain poor due to the onset of heart failure (3). Ischemic myocardial injury is increasingly recognized to orchestrate adverse cardiac tissue remodeling, resulting in ventricular stiffening, dilation, and ultimately, heart failure with reduced ejection fraction (HFrEF) (4). Sustained immune reactions represent central elements to adverse cardiac tissue remodeling (5), and thereby offer opportunities for targeted interventions to mitigate and even reverse HFrEF in response to MI (6).

Initial waves of infiltrating bone marrow-derived leukocytes, including monocytes, into ischemic cardiac tissue in response to MI have long been recognized as a source of tissue repair (7). However, exaggerated cardiac immune cell recruitment sustains inflammation that results in adverse ventricular remodeling and cardiac functional decline (8). Recent studies have identified the Type 1 Interferon (T1 IFN) response axis as a critical element responsible for adverse cardiac remodeling and HF in response to MI (9).

Cytosolic DNA sensors in phagocytic macrophages, including cyclic GMP-AMP synthase-stimulator of interferon genes (cGAS-STING), recognize double-stranded DNA (dsDNA) released as a damage- associated molecular pattern (DAMP) during ischemic cardiomyocyte cell death, a hallmark of MI (10, 11). In response, phagocytic macrophages activate the interferon regulatory factor 3 (IRF3) transcription factor, which in turn triggers the T1 IFN signaling cascade, resulting in the upregulated expression of interferon-stimulated genes (ISGs) that sustain cardiac myeloid cell recruitment, inflammation and maladaptive ventricular remodeling (9). Furthermore, the T1 IFN signaling cascade was found to be triggered within the bone marrow via sympathetic responses to MI, accentuating myelopoiesis and inflammatory myeloid cell recruitment to cardiac tissue (12, 13). Inhibiting any component of the T1 IFN axis mitigates undesirable immune reactions during cardiac tissue repair post-MI, supporting the targeting of this inflammatory signaling axis as a therapeutic strategy for HF (14).

Extracellular vesicles, including exosomes are increasingly recognized as sources of intercellular communication, owing to their capacity to transport microRNAs, proteins, metabolites, and other bioactive molecular cargo to recipient cells (15–17). Studies, including from our laboratory, have reported on macrophage-derived exosomes as mediators of immunometabolic signaling in cardiovascular inflammation and diabetes (17–19). While exosomes produced by macrophages polarized into an M1 pro-inflammatory state drive inflammatory signaling in the cardiovascular system (18), those produced by anti-inflammatory M2 macrophages suppress NF-κB signaling and foster the resolution of inflammation and atherosclerosis (19). Outcomes of our recent studies on exosomes produced by Interleukin-4 (IL-4) polarized human THP-1 macrophages (THP1-IL4-exo) highlighted their capacity to control cardiometabolic inflammation, improve insulin resistance, and normalize blood glucose levels in obese diabetic mice fed a lipid rich diet (17).

Building on these observations, we sought to test the effectiveness of THP1-IL4-exo in suppressing maladaptive cardiac remodeling in our previously described mouse model of diet-induced occlusive coronary atherosclerosis, MI and HFrEF (20, 21). Our findings uncover the capacity for THP1-IL4-exo to exert profound cardioprotection and recovery of cardiac function in mice with HFrEF caused by MI.

Mechanistically, cardioprotective properties of THP1-IL4-exo originated from their control of T1 IFN signaling in myeloid cells residing within the bone marrow, circulation, and cardiac tissue. Collectively, our findings introduce the therapeutic potential of THP1-IL4-exo, a type of M2-macrophage exosomes, in reversing cardiac functional decline and HFrEF in response to MI caused by occlusive coronary atherosclerosis.

## Materials and Methods

### Animal studies

ApoE^h/h^/SR-B1^−/−^ mice were bred to Mx1-Cre transgenic mice, and the resulting offspring were used to conduct *in vivo* studies. At 20-24 weeks of age, ApoE^h/h^/SR-B1^−/−^/Mx1-Cre^+^ mice were fed a Paigen’s Atherogenic Rodent Diet (Research Diets) for 4.5 weeks. This time point was chosen to induce occlusive coronary atherosclerosis, MI, and CHF in the mice. Afterwards, 250 µg of polyinosinic:polycytidylic acid (pIpC) was injected into each mouse over the course of two days, and they were also switched to a chow diet (Teklad). pIpC-mediated repair of ApoE resulted in lipid lowering. During this time, cohorts of mice were also injected with either THP1-IL4-exo or PBS control. This treatment paradigm was continued for 4.5 weeks. Data collection and analyses were conducted in a blinded fashion. All mice were housed and bred in specific pathogen-free conditions in the Animal Research Facility at the San Francisco Veterans Affairs Medical Center or at UCSF Mission Bay. All animal experiments were approved by the Institutional Animal Care and Use Committee at the San Francisco Veterans Affairs Medical Center.

### Echocardiographic analysis of heart function

Transthoracic echocardiography was performed with a Vevo F2 (Fujifilm Visualsonics) system using the UHF46x 46-20 MHz transducer. Mice were lightly sedated with isoflurane, shaved, secured to the heating platform in a supine position, and monitored for body temperature and consistent heart rate. Ultrasound gel (Aquasonic) was applied to the shaved chest. Two-dimensional (2D) long-axis images of the LV were recorded at the plane of the aortic and mitral valves, where the LV cavity is largest. Additionally, 2D short-axis images were obtained at the papillary muscle level. All measurements were made from digital images capture on cine loops at Time points A, B, and C. The Vevo Lab analysis software (Fujifilm Visualsonics) was used to analyze data for EF, FS, LVESV, LVEDV, and SV.

### Cell culture

The human monocytic cell line Tohoku Hospital Pediatrics-1 (THP-1) was purchased from the University of California, San Francisco (UCSF) Cell and Genome Engineering Core as an authenticated stock. Cells were cultured in RPMI 1640 media (Corning) supplemented with 10% fetal bovine serum (Gibco), 1% GlutaMAX (Gibco), and 1% penicillin-streptomycin (Gibco, USA). THP-1 cells were grown and expanded in a T-75 flask (Thermo Fisher Scientific) until they reached a density of 10^6^ cells/mL. Once confluent, cells were seeded into 100mm plates (VWR, USA) at 4 × 10^6^ cells/plate. Phorbol 12-myristate 13-acetate (PMA; Thermo Fisher Scientific) was added to the plates at a concentration of 25 ng/mL for 48 hours, differentiating the THP-1 cells into macrophages. THP-1 macrophages were then cultured in PMA-free media for an additional 48 hours, followed by culturing in exosome-free media (EFM) supplemented with 20 ng/mL of recombinant human IL-4 (Peprotech) to produce IL-4 conditioned media (19).

### THP-1 macrophage exosome isolation and nanoparticle tracking analysis

Exosome isolation and characterization experiments were performed in accordance to the MISEV 2023 guidelines (42). After growing the differentiated THP-1 macrophages in dishes with PMA-free media for 48 hours, the media was removed, the plates were washed with PBS (Corning) twice, and EFM supplemented with IL-4 was added for 24 hours. The CM was collected the following day and centrifuged at 400 ×g for 10 mins, poured into a new tube and centrifuged at 2,000 × g for 20 mins, and filtered with a 0.2 μm membrane. THP1-IL4-exo were then isolated using cushioned-density gradient ultracentrifugation (C-DGUC). The filtered CM was first centrifuged on a 60% OptiPrep cushion (StemCell Technologies) at 100,000 × g for 3 hours and at 4 °C (Type 45 Ti, Beckman Coulter) to concentrate the exosomes. Afterwards, the cushion was collected and centrifuged along with a 5%, 10%, and 20% OptiPrep density gradient at 100,000 × g overnight and at 4 °C (SW 40 Ti, Beckman Coulter). The following day, twelve 1 mL fractions were collected, starting from the top of the Beckman tube. Fraction 7 from the density gradient was dialyzed in PBS using the Slide-A-Lyzer MINI Dialysis Device (Thermo Fisher Scientific) and used for characterization, *in vitro*, and *in vivo* experiments.

The particles in Fraction 7 were subjected to Nanoparticle Tracking Analysis (NTA) using the NanoSight LM10 (Malvern Panalytical), which provided size and concentration data. Samples were diluted in 1:100 PBS and measured in triplicate. The analysis settings were optimized and kept identical for each sample, and data were analyzed using the NanoSight NTA v3.3 software. All exosome samples were stored at 4 °C and used after isolation or stored at -80 °C.

### Labeling and in vitro tracking of THP-1 macrophage exosome biodistribution

THP1-IL4-exos were labeled with 1,1’-dioctadecyl-3,3,3’3’ tetramethylindotricarbocyanine iodide (DiR [DilC18(7)]; Invitrogen). The dye was added to the 60% OptiPrep cushion at a concentration of 10 μM, incubated for 20 minutes, then isolated following the same C-DGUC protocol as previously stated. After dialysis and NTA characterization, 10^10^ particles of DiR-labeled THP1-IL4-exo or PBS were injected into ApoE^h/h^/SR-B1^−/−^/Mx1-Cre^+^ mice that had been fed a Paigen’s diet for 4.5 weeks, induced with pIpC, and returned to a chow diet. After 6 hours, the mice were sacrificed, perfused with PBS, and their blood, tibias, femurs, and hearts were collected. The DiR fluorescence signal was imaged using the Odyssey CLx Infrared Imaging System and analyzed using the Image Studio software (LI-COR Bio).

### Transmission electron microscopy

Exosome morphology was assessed by electron microscopy by loading 7 × 10^8^ exosomes onto a glow-discharged 400-mesh Formvar-coated copper grid (Electron Microscopy Sciences). The particles were left to settle for 2 min, after which the grids were washed four times with 1% uranyl acetate. Excess uranyl acetate was blotted off gently with filter paper. Grids were then dried and imaged at 120 kV using a Tecnai 12 transmission electron microscope (FEI).

### Transcriptional profiling of gene expression in circulating Ly-6C^hi^ monocytes

Paigen diet-fed ApoE^h/h^/SR-B1^−/−^/Mx1-Cre^+^ mice that were infused three times a week with THP1-IL4-exo vs PBS ctrl intraperitoneally (IP) were anesthetized with isoflurane (Baxter), after which their blood was collected via retro-orbital puncture with heparinized micro-hematocrit capillary tubes (Fisher Scientific) into tubes with 0.5M EDTA (Invitrogen). Red blood cells were then lysed with 1x RBC Lysis Buffer (BioLegend). Leukocytes were then stained with anti-CD45 (Clone 30-F11), -CD11b (Clone M1/70), - CD115 (Clone AFS98), and -Ly-6C (Clone HK1.4, all from BioLegend) for 30 min at 4°C and in the dark. The antibody dilutions ranged from 1:100-1:200. Ly-6C^hi^ monocytes were sorted using the BD FACSAria Fusion (BD Biosciences). RNA was isolated from the sorted Ly-6C^hi^ monocytes using the RNeasy Mini Kit (QIAGEN), quantified using the Quant-iT RiboGreen RNA Assay Kit, and 10 ng of RNA was amplified using the nCounter Low RNA Input Amplification Kit before expression analysis with the nCounter analysis system (NanoString Technologies). The mRNA detection procedure was performed according to the manufacturer’s instructions using the autoimmune profiling panel for mice. The number of mRNA molecules counted was imported into the NSolver 4.0 (NanoString Technologies) with default settings, corrected and normalized against the reference genes annotated in the kit that were found to be stable. The software generated an Advanced Analysis Report, and results are considered statistically significant when the p value is ≤ 0.05.

### Isolation of CD11b^+^ myeloid cells from mouse tissues

ApoE^h/h^/SR-B1^−/−^/Mx1-Cre^+^ were sacrificed and perfused with PBS. Afterwards, their tibias, femurs, and hearts were collected. Tibias and femurs were flushed by centrifugation at 3,000 x g for 3 mins to obtain the cells of the bone marrow. Hearts were digested using the Multi Tissue Dissociation kit (Miltenyi Biotec) and following the manufacturer’s instructions. Both cell suspensions were then subjected to 1x RBC Lysis Buffer (BioLegend), followed by isolation of CD11b cells using mouse CD11b MicroBeads (Miltenyi Biotec), which was performed according to the manufacturer’s instructions. Cells were then placed in QIAzol Lysis Reagent (QIAGEN) for subsequent RNA extraction.

### RNA extraction and gene expression analysis using qRT-PCR

For heart tissues that were not subjected to CD11b MicroBead capturing, they were homogenized in QIAzol with a Tissue-Tearor (BioSpec). Total RNA from cells in QIAzol was then extracted using the RNeasy Mini Kit (QIAGEN) according to the manufacturer’s protocol. RNA was then quantified using NanoDrop and reverse transcribed using the iScript Reverse Transcription Supermix (Bio-Rad). qPCR reactions were performed using the iTaq Universal SYBR Green Supermix (Bio-Rad) and run on the CFX Opus 384 Real-Time PCR System (Bio-Rad). Cycle threshold (Ct) values were normalized to the housekeeping genes *B2m* and *Gapdh* and analyzed using the Bio-Rad CFX Maestro software (Bio-Rad). All reactions were performed in triplicate.

### Circulating and tissue-associated leukocyte detection using flow cytometry

Blood was collected, processed, and stained as previously described. Ly-6C^hi^ monocytes were subjected to flow cytometric analysis instead of cell sorting.

For the bone marrow, cells were flushed and collected as previously described. Cells were washed, re-centrifuged, and stained in FACS buffer containing a lineage-marker mix of biotinylated anti-CD4 (Clone RM4.5), -CD8 (Clone 53-6.7), -CD45R/B220 (Clone RA3-6B2), -TER-119 (Clone TER-119), -Gr-1 (Clone RB6-8C5), and -CD127 (Clone A7R34, all from BioLegend) for 30 min at 4°C and in the dark. Cells were then stained with anti-CD34 (Clone RAM34), -CD48 (Clone HM48.1), -Ly-6A/E/Sca-1 (Clone D7), -CD135 (Clone A2F10), -CD117/c-Kit (Clone 2B8, all from Invitrogen eBioscience), -CD150/SLAM (Clone TC15-12F12.2), - CD16/32 (Clone 93), -CD41 (Clone MWReg30, all from BioLegend), and Streptavidin (Clone 563858, BD Biosciences) for 90 min at 4°C and in the dark. Cells were gently mixed every 20 min. The antibody dilutions ranged from 1:100-1:200.

For the heart, tissues were digested using the Multi Tissue Dissociation kit (Miltenyi) and following the manufacturer’s instructions. Red blood cells were then lysed with 1x RBC Lysis Buffer (BioLegend).

Nonspecific binding was blocked using TruStain FcX anti-mouse CD16/32 antibodies (BioLegend) for 10 min at 4°C in FACS buffer, followed by staining with anti-CD45 (Clone 30-F11), -CD11b (Clone M1/70), -CD64 (Clone X54-5/7.1), -Ly-6C (Clone HK1.4), -Ly-6G (Clone HK1.4), -CD119/CCR2 (Clone SA20361.1), and -TIM-4 (Clone RMT4-54, all from BioLegend) for 30 min at 4°C and in the dark. The antibody dilutions ranged from 1:100-1:200.

All euthanized mice were fully perfused with PBS before the tissues were harvested. All flow cytometric analyses were conducted using a CytoFLEX S Flow Cytometer (Beckman Coulter). All flow cytometry data were collected and analyzed using FlowJo v10.9.0 (FlowJo).

### Measurement of cholesterol levels in mouse plasma

Peripheral blood was collected as indicated above and spun at 4,000 rpm for 30 min at 4 °C to collect the plasma. Plasma cholesterol was measured using the Cholesterol E Assay Kit (Fujifilm Wako Pure Chemical Corporation).

### Quantification and statistical analysis

Statistical analyses were performed using GraphPad Prism v8, using the unpaired, two-tailed, Student’s t test (two groups). ∗p < 0.05, ∗∗p < 0.01, ∗∗∗p < 0.001. All error bars represent the mean ± the standard error of the mean (SEM). All experiments were repeated at least twice or performed with independent samples.

## Results

### Cardioprotective properties of M2-polarized human macrophage exosomes in diet-induced occlusive coronary atherosclerosis, MI, and HF

In this study, male ApoE^h/h^/SR-B1^−/−^/Mx1-Cre^+^ mice were fed an atherogenic Paigen’s diet for 4.5 weeks to induce hyperlipidemia, occlusive coronary atherosclerosis, and MI as we previously reported (20, 21). (**Figure 1A**). At that time point (Time point B), mice were switched to chow diet and intra-peritoneally (IP) injected twice over two days with 250 µg of polyinosinic:polycytidylic acid (pIpC) to activate the Mx1- Cre recombinase that repairs the hypomorphic ApoE^h/h^ (HypoE) alleles, restoring physiological ApoE expression in the liver that rapidly normalizes plasma lipid levels as we previously reported (21, 22).

**Fig 1.**
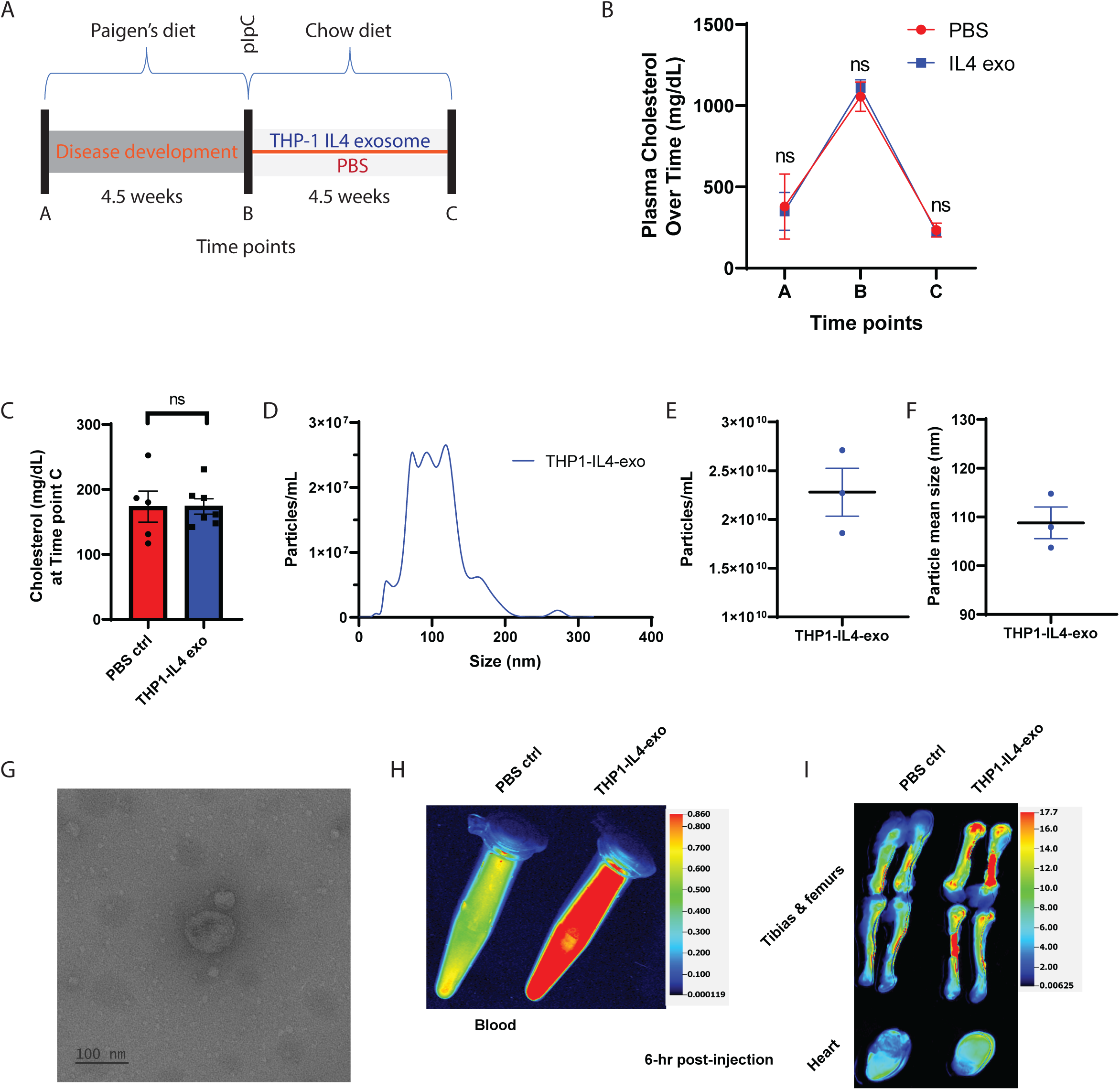
Study design utilizing ApoE^h/h^/SR-B1^−/−^Mx1-Cre^+^ mice and biophysical parameters of THP1- IL4-exo. (A) Schematic depicting the time course involving diet modification and treatment with exosomes. (B) Plasma cholesterol levels in both treatment groups prior to starting Paigen’s diet (Time point A), after 4.5 weeks on the diet (Time point B), and after 4.5 weeks of THP1-IL4-exo treatment or PBS (Time point C). (C) Plasma cholesterol levels at Time point C; n = 5-7 in each group, pooled from two independent experiments. Data are represented as mean ± SEM. (D) Representative concentrations and size distributions of THP1-IL4-exo purified from THP-1 conditioned cell culture supernatant after a 24 hr culture period as determined using Nanoparticle Tracking Analysis (NTA). (E and F) Average particle concentration per milliliter (E) and mean diameter in nanometers (F) (n = 3). (G) Electron micrograph of purified THP1-IL4-exo. Scale bar: 100nm. (H and I) Images of DiR fluorescence in the blood (H), tibias, femurs, and hearts (I) 6 hr post-injection from ApoE^h/h^/SR-B1^−/−^Mx1-Cre^+^ mice that had been fed a Paigen’s diet for 4.5 weeks, induced with pIpC, returned to a chow diet, and injected IP with 10^10^ particles of DiR-labeled THP1-IL4-exo or PBS control.

Subsequently, mice were left to recover from MI while undergoing tri-weekly IP treatments with 10^10^ particles of THP1-IL4-exo or saline (**Figure 1A**). At time points A, B, and C, blood was collected to determine cholesterol levels and the numbers of monocytes and neutrophils, while high-resolution ultrasound served to record parameters of cardiac function. **Figure 1B** shows the efficacy of Cre- mediated gene repair of the HypoE alleles and chow diet restoration in normalizing plasma cholesterol levels at Time points C. **Figure 1C** further highlights the normalized plasma cholesterol levels among both treatment groups of mice at the time of tissue collection (n = 5-7).

THP1-IL4-exo used in this study were produced and isolated via Cushioned-Density Gradient Ultracentrifugation (C-DGUC) (23, 24) and confirmed to have similar concentrations, size distributions, and morphologies (**Figure 1D-G**) as reported in our prior study (17). To assess their biodistribution in this model, freshly prepared THP1-IL4-exo were labeled with 1,1’-dioctadecyl-3,3,3’3’ tetramethylindotricarbocyanine iodide (DiR [DilC18(7)]) and injected IP into ApoE^h/h^/SR-B1^−/−^/Mx1-Cre^+^ that had been fed a Paigen’s diet for 4.5 weeks, induced with pIpC, and returned to a chow diet. The mice were given one injection of 10^10^ particles of DiR-labeled THP1-IL4-exo or PBS. Six hours post treatment, the presence of DiR-positive exosomes were detected in the blood, tibias, femurs, and heart of the mice (**Figure 1H-I**) as we reported in the study of obese diabetic mice (17).

### THP1-IL4-exo treatments improve survival and left ventricle cardiac function reversing HFrEF post-MI

To begin investigating cardioprotective properties of THP1-IL4-exo, we tested their impact in prolonging longevity and cardiac function post-MI. As shown in **Figure 2A**, THP1-IL4-exo treatments led to a significant improvement in survival rates up to 20 weeks post MI. Next, we performed serial echocardiographic measurements of left ventricular (LV) function, including ejection fraction (EF), fractional shortening (FS), left-ventricular end-systolic volume (LVESV), left-ventricular end-diastolic volume (LVEDV), and stroke volume (SV) in mice of both treatment groups at all three Time points (A, B, C). Representative echocardiographic images of the short (left) and long axes (middle) of the heart are shown in **Figure 2B**, including a representative image of the quantitative analysis using the axes images (right). As shown in **Figure 2C**, all mice experienced an expected decline in LV function during Time points A and B, corresponding to the period of Paigen diet consumption, which results in occlusive coronary atherosclerosis, MI, and onset of HF with reduced ejection fraction (HFrEF) as reported in our prior studies of the model (20, 21). Following HypoE allele repair and diet change at Time point B, tri- weekly IP injections of THP1-IL4-exo led to significantly improved LV function in mice until Time point C, where numerous parameters of cardiac function were detected to return to baseline levels prior to diet initiation (**Figure 2C**). Data in **Figure 2D** show improved parameters of LV function (EF, FS, LVESV, LVEDV, SV) among groups of mice at Time point C, following 4.5 weeks of treatment with THP1-IL4-exo, highlighting reversal of LV functional decline compared to mice treated with sham, which all experienced further cardiac functional decline.

**Fig. 2.**
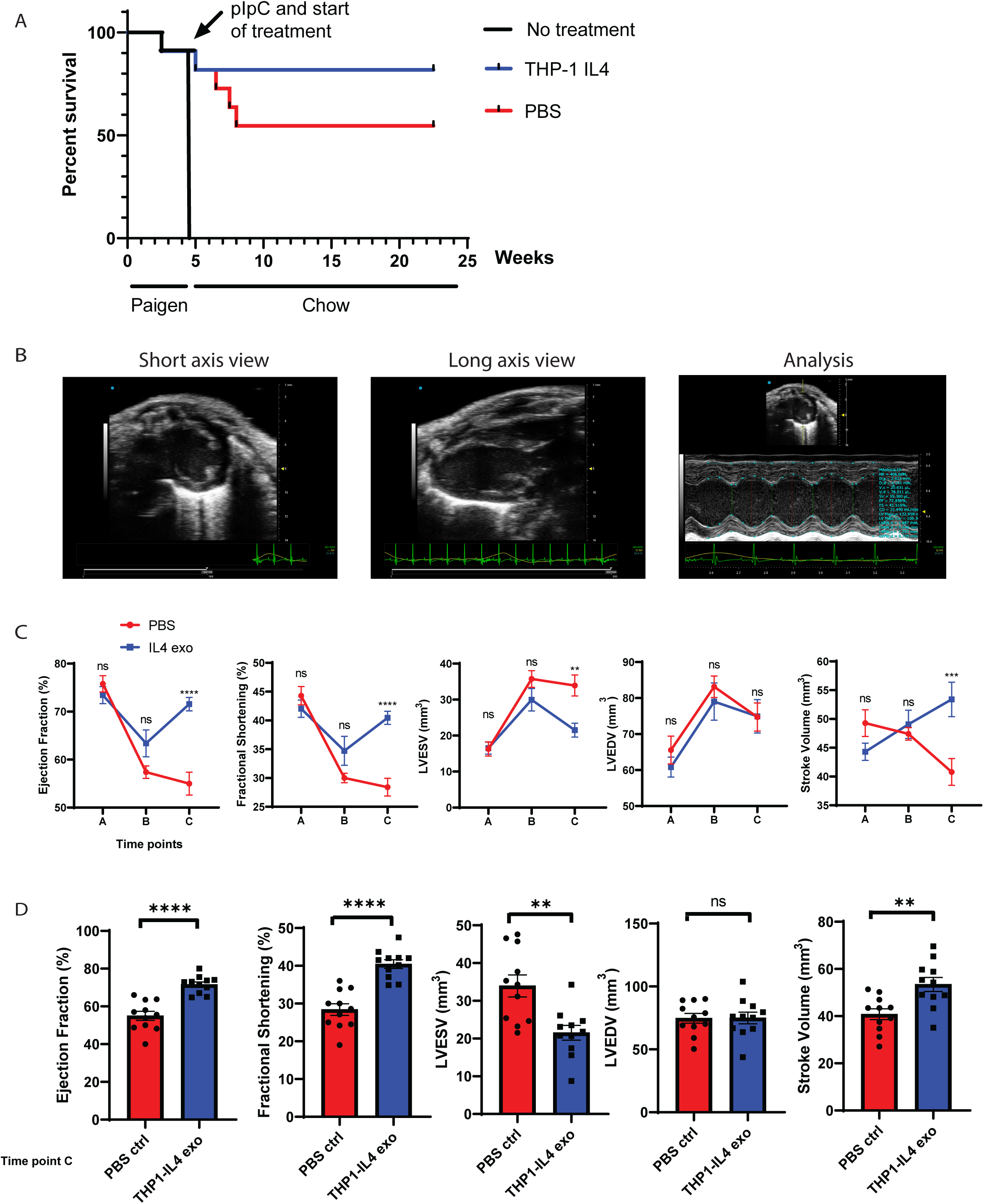
THP1-IL4-exo improves survival and left ventricle function in mice with diet-induced myocardial infarction & heart failure. (A) Kaplan-Meier survival curve ApoE^h/h^/SR-B1^−/−^Mx1-Cre^+^ mice fed Paigen diet followed by a pIpC injection and chow diet, treated with THP1-IL4-exo or PBS for up to 23 weeks. p < 0.06 as determined by a log-rank test (n = 10 in each group). (B) Representative echocardiographic images of the short axis view (left), the long axis view (middle), and the quantitative data analysis of the axes images (right). (C) Echocardiographic measures of left ventricular performance over the three Time points, including ejection fraction (EF), fractional shortening (FS), left-ventricular end systolic volume (LVESV), left- ventricular end diastolic volume (LVEDV), and stroke volume (SV). D) EF, FS, LVESV, LVEDV, and SV measured at Time point C; n = 10-11 in each group, data pooled from two independent experiments. *p < 0.05, **p < 0.01, ***p < 0.001, and ****p < 0.0001 as determined using unpaired Student’s t test. Data are represented as mean ± SEM.

### THP1-IL4-exo treatments control the numbers of Ly-6C^hi^ monocytes and neutrophils in the circulation

Next, we examined the number of myeloid cells in the circulation, recognized for their propensity to traffic into the heart following MI (7, 8, 13, 25). Blood collected from both groups of mice at Time points A, B, and C was examined using flow cytometric analysis. **Figure 3A** illustrates the gating strategy used to collect data on the numbers of circulating Ly-6C^lo^ and Ly-6C^hi^ monocytes, as well as neutrophils. No significant differences were observed among the three groups of myeloid cells in the circulation between Time points A and B. However, significant reductions in the numbers of circulating Ly-6C^hi^ and neutrophils were apparent in THP1-IL4-exo-treated mice at Time point C, which also tended to display higher numbers of circulating Ly-6C^lo^ cells (**Figure 3B**). These data further support the benefits of systemic immunosuppression offered by the THP1-IL4-exo in this mouse model of cardiovascular inflammation where MI is a dominant stimulant of myelopoiesis (7, 8, 13, 25).

**Fig. 3.**
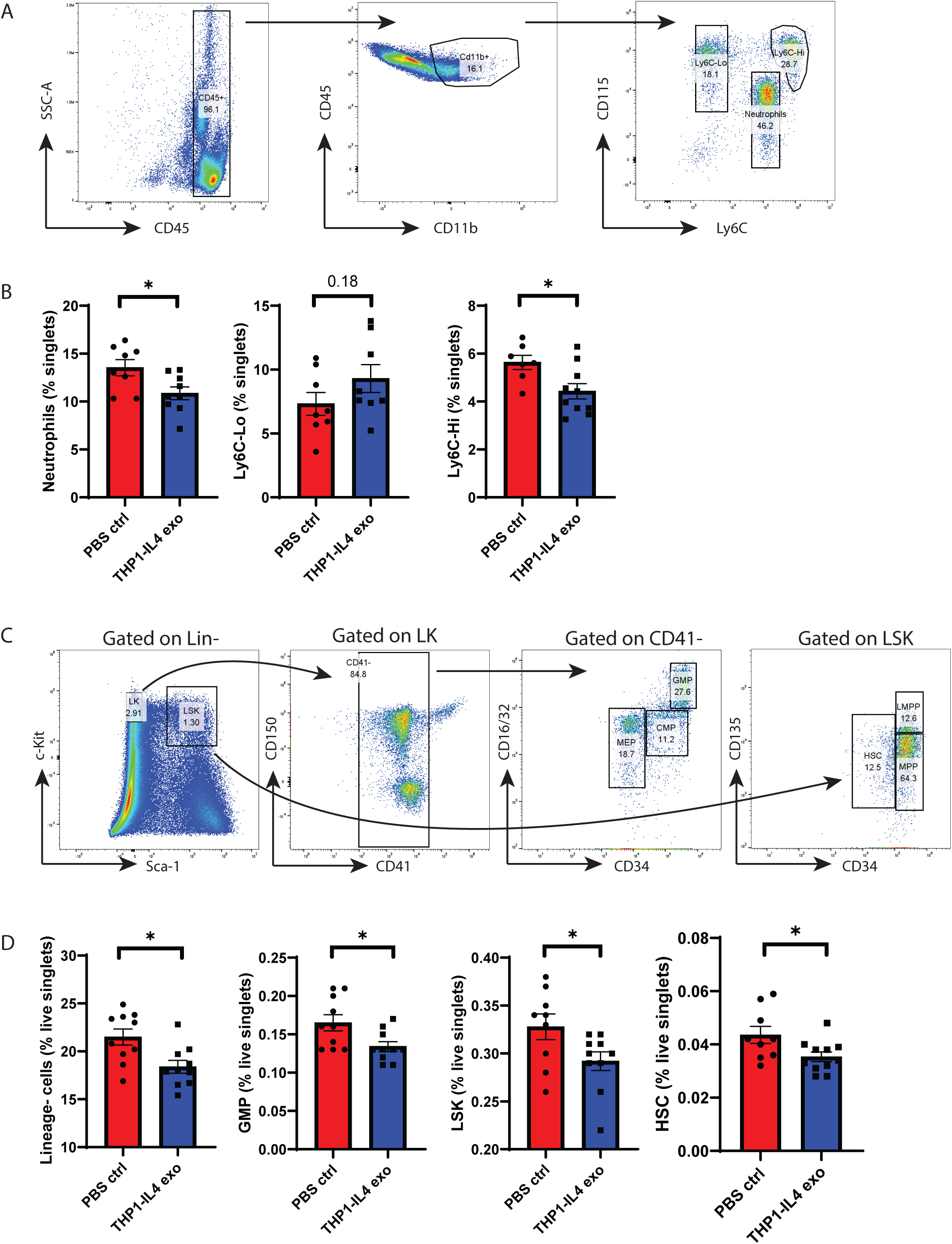
THP1-IL4-exo control the number of myeloid cells in the circulation while suppressing hematopoiesis and myelopoiesis in the bone marrow. (A) Representative flow cytometry plots of leukocyte subsets from peripheral blood. (B) Quantification of neutrophils, Ly-6C^lo^, and Ly-6C^hi^ populations from the peripheral blood of THP1-IL4- exo vs PBS ctrl mice. (C) Representative flow cytometry plots of leukocyte progenitor subsets from the bone marrow. (D) Quantification of Lineage negative (Lin^−^), granulocyte-monocyte progenitor (GMP), Lin^−^/Sca1^+^/c-Kit^+^ (LSK), and hematopoietic stem cell (HSC) populations from the bone marrow of mice treated with THP1- IL4-exo vs PBS ctrl; n = 6-11 in each group, data pooled from two independent experiments. *p < 0.05, as determined using unpaired Student’s t test. Data are represented as mean ± SEM.

### THP1-IL4-exo treatments suppress hematopoiesis and myelopoiesis in the bone marrow

To further explore benefits of THP1-IL4-exo treatment in suppressing the number of inflammatory myeloid cells in the circulation, we examined populations of leukocyte progenitor cells in the bone marrow (BM). **Figure 3C** shows the gating strategy used to detect Lineage negative (Lin^-^) cells, granulocyte-monocyte progenitor (GMP) cells, Lineage^-^/Sca-1^+^/c-Kit^+^ (26) cells, and hematopoietic stem cells (HSCs). All mice treated with THP1-IL4-exo displayed reduced populations of Lin^-^ cells, GMP cells, LSK cells, and HSCs (**Figure 3D**). Reduced hematopoiesis and myelopoiesis highlight the immunosuppressive properties of THP1-IL4-exo within the BM of mice that experienced MI, which mirrors our prior findings observed among obese diabetic mice with hyperlipidemia (17, 19).

### THP1-IL4-exo treatments suppress the Type 1 Interferon pathway and inflammation in circulating Ly-6C^hi^ monocytes

Because Ly-6C^hi^ monocytes are widely recognized for contributing to HF by driving inflammation, fibrosis, and adverse LV remodeling following MI (27), we sought to investigate their transcriptional responses to four weeks of THP1-IL4-exo treatments. To do so, circulating Ly-6C^hi^ monocytes from both groups of mice were isolated from freshly-drawn blood using a FACS sorter, and their extracted RNA was processed for unbiased transcriptional profiling of genes involved in inflammation and metabolism using a NanoString platform. Volcano plots shown in **Figure 4A** highlight the differential expression of genes among Ly-6C^hi^ monocytes collected from both groups of mice. Most notably, repeated THP1-IL4-exo treatments led to a downregulation of inflammatory genes that included: *Socs1*, *Flt1*, *Jun*, *Irf4*, *Tlr9*, *Txnip*, *Tbk1*, *Traf6*, and *Irak1*. The heatmap of differentially expressed genes shown in **Figure 4B** highlights a reduction of numerous genes associated with the T1 IFN pathway. Together, the panoply of inflammatory genes controlled by THP1-IL4-exo supports their capacity to suppress inflammatory activities, including those associated with the T1 IFN axis, in circulating Ly-6C^hi^ monocytes.

**Fig. 4.**
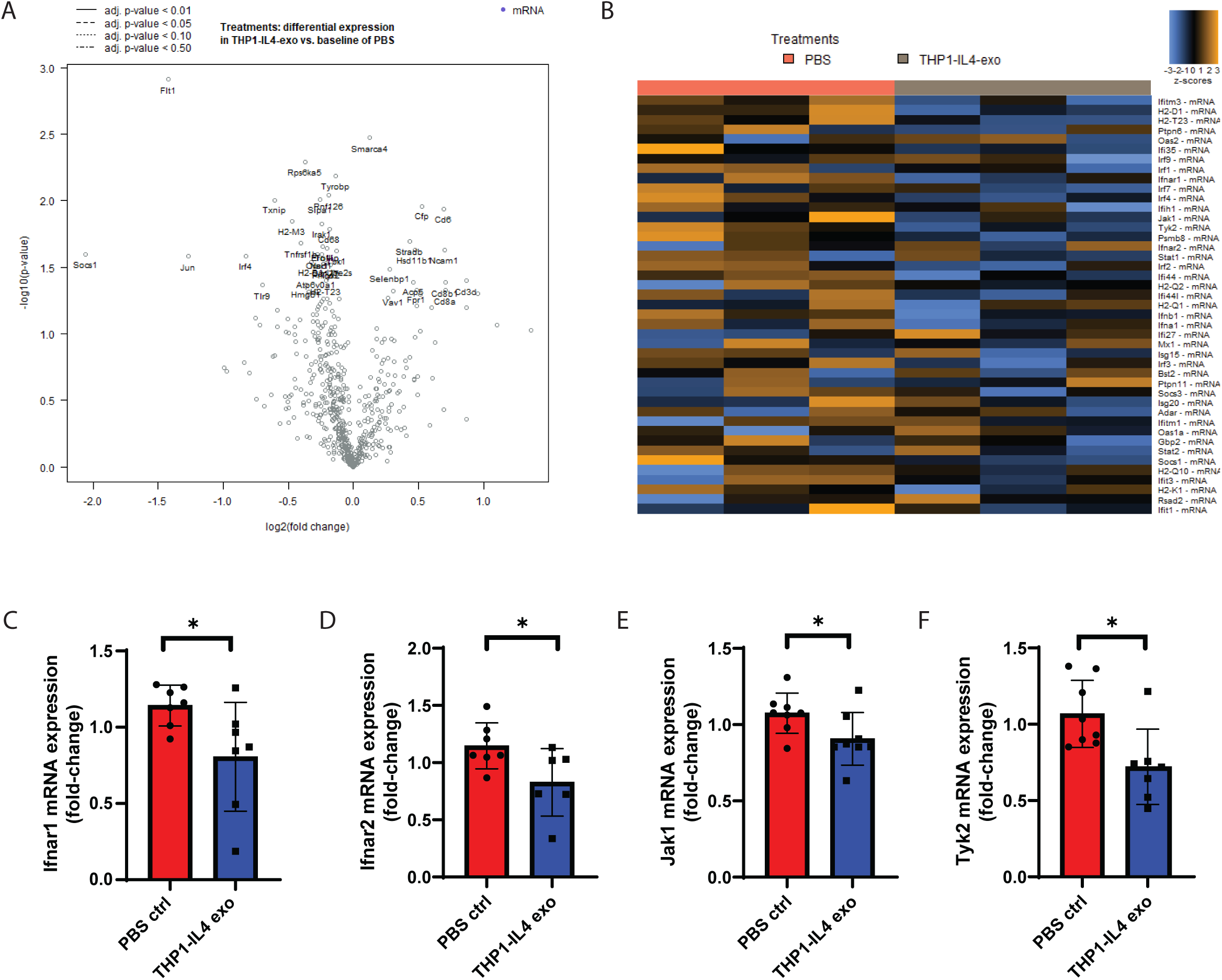
THP1-IL4-exo suppress the Type 1 Interferon response in circulating Ly-6C^hi^ monocytes and bone marrow CD11b^+^ myeloid cells. (A and B) NanoString transcriptomic analysis of Ly6C^hi^ monocytes sorted from the peripheral blood of ApoE^h/h^/SR-B1^−/−^Mx1-Cre^+^ mice. The volcano plot depicts the most significant gene changes found in the Ly-6C^hi^ monocytes (A). The heatmap illustrates the differential expression of genes of the T1 IFN pathway (B) (n = 3). (C-F) qRT-PCR analysis of *Ifnar1* (C), *Ifnar2* (D), *Jak1* (E), and *Tyk2* (F) mRNA expression in bone marrow-derived CD11b^+^ myeloid cells from mice treated with THP1-IL4-exo vs PBS ctrl. Gene expression is normalized to *B2m* and *Gapdh* mRNA; n = 6-8, data pooled from two independent experiments. *p < 0.05, as determined using unpaired Student’s t test. Data are represented as mean ± SEM.

### THP1-IL4-exo treatments suppress the Type 1 Interferon pathway in bone marrow-derived myeloid cells

In light of transcriptomic data supporting robust reprogramming of inflammatory pathways in Ly-6C^hi^ monocytes exposed to THP1-IL4-exo, we next sought to investigate whether the T1 IFN response was similarly modulated in myeloid cells residing within the BM, recognized as a source of inflammatory activity following MI (12). We did so by isolation of CD11b^+^ myeloid cells from the tibias and femurs of both groups of mice. Next, RNA was extracted and subjected to RT-qPCR analysis. Data shown in **Figure 4C** depict a significant downregulation in the expression of genes encoding the two subunits of the IFNAR, the receptor recognized as central to the activation of T1 IFN signaling (9). Additionally, *Jak1* and *Tyk2*, genes encoding for kinases responsible for signal transduction following IFNAR activation (28), were also significantly downregulated. Together, these data highlight the efficacy of THP1-IL4-exo treatments in suppressing T1 IFN signaling within myeloid cells residing in the BM of mice that experience MI.

### THP1-IL4-exo treatments suppress cardiac myeloid cell recruitment while preserving populations of resident cardiac macrophages

To investigate mechanisms through which treatments with THP1-IL4-exo controlled HFrEF, we tested their impact on modulating cardiac inflammation post-MI. We did so by performing flow cytometric analyses of leukocyte populations in enzymatically digested cardiac tissues of mice euthanized at Time point C. Using the gating strategy shown in **Figure 5A**, we detected reduced numbers of monocytes, Ly- 6C^+^/CD64^-^ infiltrating monocytes, and neutrophils in the cardiac tissue of THP1-IL4-exo treated mice (**Figure 5B**). Interestingly, THP1-IL4-exo treatments appeared to foster an augmentation in the number of resident cardiac macrophages (rcMacs), identified by the expression of the cell surface marker T-cell membrane protein 4 (TIM-4^+^/CCR2^-^) (**Figure 5C**) (26). Furthermore, recruited M2-like monocyte-derived macrophages positive for TIM-4 and CCR2 expression (TIM-4^+^/CCR2^+^) were found to be more abundant in the cardiac tissue of mice treated with THP1-IL4-exo (**Figure 5C**). Together, these data demonstrate the efficacy of THP1-IL4-exo treatments in suppressing inflammation while augmenting the number of anti- inflammatory tissue-repair monocyte-derived and resident macrophages in the cardiac tissue of mice that develop HFrEF post-MI.

**Fig. 5.**
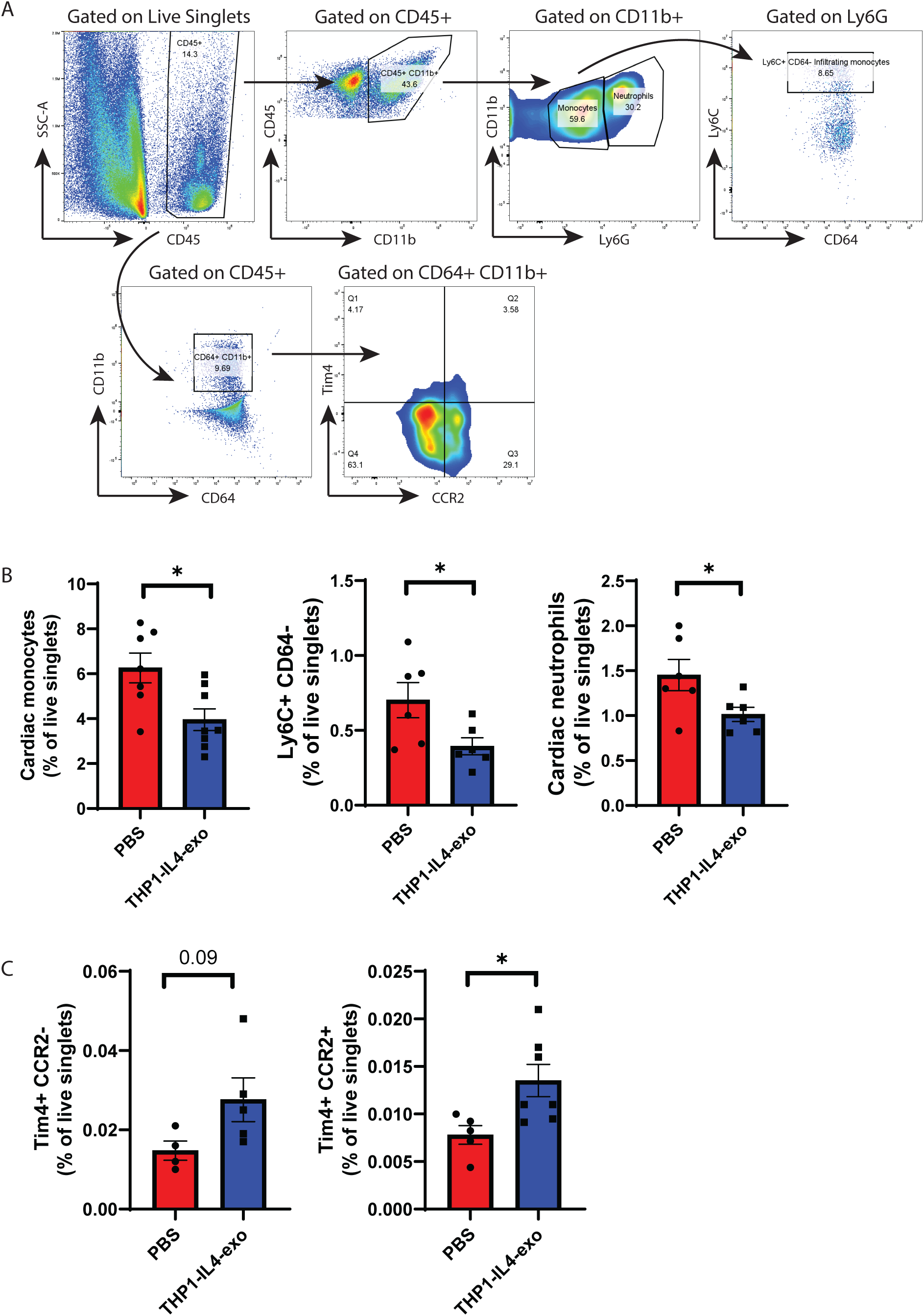
THP1-IL4-exo suppress inflammatory myeloid cell recruitment into cardiac tissue and preserve resident macrophages. (A) Representative flow cytometry plots of leukocyte subsets from digested whole heart tissues. (B) Quantification of cardiac monocytes, Ly-6C^+^/CD64^−^ infiltrating monocytes, and neutrophils from the digested hearts of mice treated with THP1-IL4-exo vs PBS ctrl. (C) Quantification of TIM-4^+^/CCR2^−^ and TIM-4^+^/CCR2^+^ macrophage populations from the digested hearts of mice treated with THP1-IL4-exo vs PBS ctrl; n = 6-7 in each group, pooled from two independent experiments. *p < 0.05, as determined using unpaired Student’s t test. Data are represented as mean ± SEM.

### THP1-IL4-exo treatments suppress Type 1 Interferon signaling in intracardiac myeloid cells that limits cardiac inflammation and adverse ventricular remodeling

Having demonstrated the potency of THP1-IL4-exo in suppressing T1 IFN activity in myeloid cells within the BM and the circulation of mice that experienced MI, we next tested their ability to target inflammation in cardiac tissue. We did so as our biodistribution study revealed the capacity for THP1-IL4- exo to traffic to infarcted heart tissue (**Figure 1F**). Thus, we investigated T1 IFN responses through an assessment of ISGs in intracardiac myeloid cells. Enzymatically digested hearts from both groups of mice at Time point C were subjected to immunomagnetic isolation of CD11b^+^ cells and extracted RNA was subjected to RT-qPCR analysis. Despite a lack of statistically significant changes in the expression levels of *Ifnar1*, *Ifnar2*, *Jak1*, and *Tyk2*, their expression levels nonetheless tended to be downregulated in cells isolated from the cardiac tissue of mice treated with THP1-IL4-exo (**Figure 6A**). However, *Mmp9*, a gene encoding for a matrix metalloproteinase recognized in driving adverse ventricular remodeling post-MI (29, 30), was significantly downregulated in intracardiac CD11b^+^ myeloid cells (**Figure 6A**). Furthermore, THP1-IL4-exo significantly downregulated the expression levels of *Nrf2*, a critical negative regulator of T1 IFN activity, supporting reduced inflammatory activity in intracardiac myeloid cells (12) (**Figure 6A**).

**Fig. 6.**
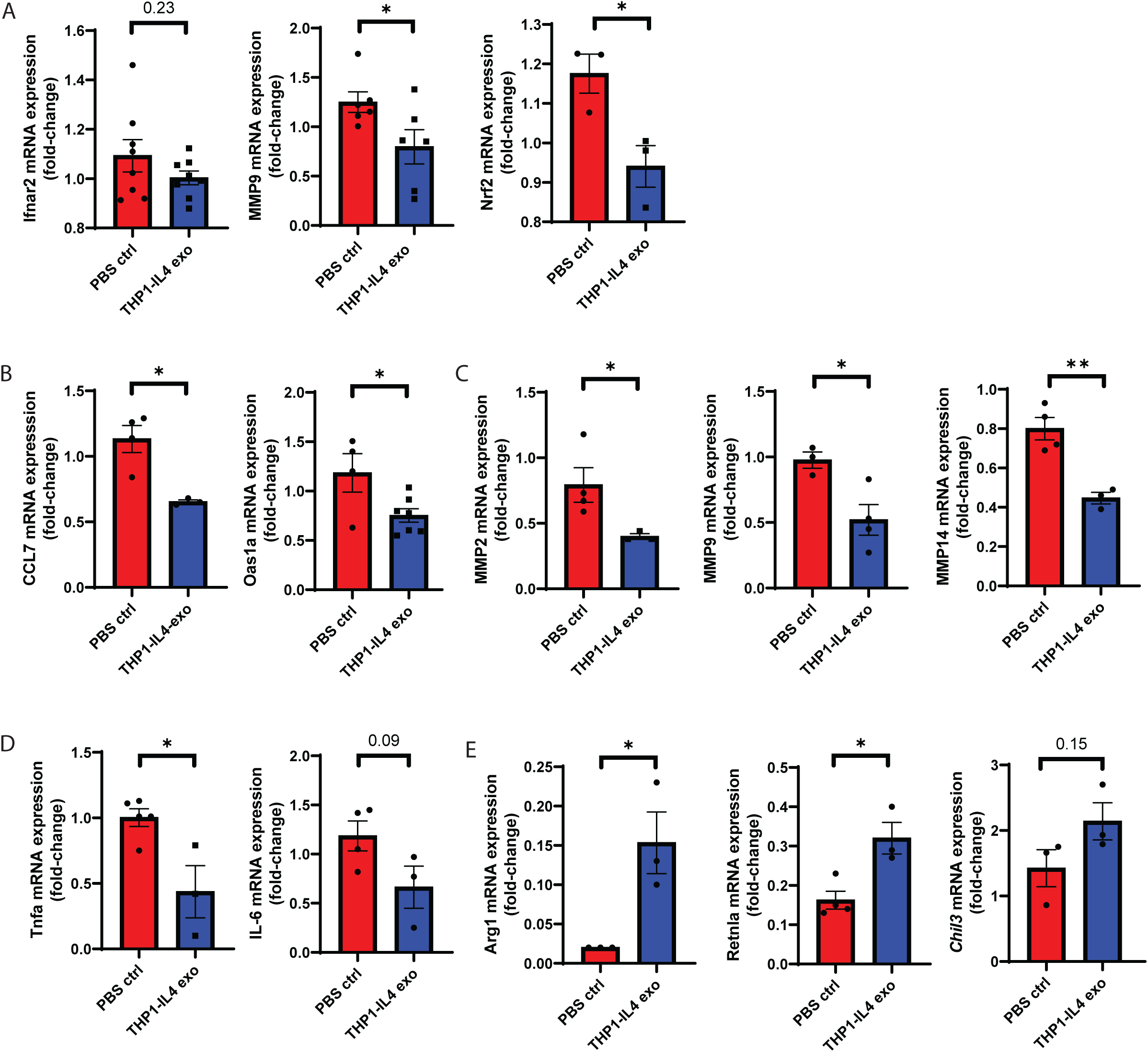
THP1-IL4-exo suppress the Type 1 Interferon response in cardiac CD11b^+^ myeloid cells and inflammation throughout cardiac tissue. (A) qRT-PCR analysis of *Ifnar2*, *Mmp9*, and *Nrf2* mRNA expression in cardiac-derived CD11b^+^ myeloid cells from mice treated with THP1-IL4-exo vs PBS ctrl. Gene expression is normalized to *B2m* and *Gapdh* mRNA; n = 3-9, data collected from one experiment or pooled from two independent experiments. (B-E) qRT-CPR analysis of *Ccl7* and *Oas1a* ISGs (B), *Mmp2*, *Mmp9*, and *Mmp14* (C), *Tnf-α* and *Il-6* (D), and *Arg1*, *Retnla*, and *Chil3* (E) mRNA expression in cardiac tissue from mice treated with THP1-IL4-exo vs PBS ctrl. Gene expression is normalized to *B2m* and *Gapdh* mRNA; n = 3-9, data collected form one experiment or pooled from two experiments. *p < 0.05, as determined using unpaired Student’s t test. Data are represented as mean ± SEM.

### THP1-IL4-exo treatments suppress the expression of inflammatory and tissue remodeling genes while enhancing the expression of those involved in anti-inflammatory activity in cardiac tissue

Lastly, we explored the capacity for THP1-IL4-exo to suppress inflammation in whole cardiac tissue. Following enzymatic digestion of cardiac tissue and extraction of RNA for RT-qPCR analysis, immune suppression by THP1-IL4-exo was evidenced by the reduced expression levels of numerous ISG genes that included Chemokine (C-C motif) ligand 7 (*Ccl7*) and 2’-5’-Oligoadenylate Synthetase 1 (*Oas1a*) (**Figure 6B**). Significant reductions were also observed for the expression of matrix metalloproteinases *Mmp2*, *Mmp9*, and *Mmp14*, recognized for contributing to adverse cardiac remodeling and ventricular dilation (**Figure 6C**) (29, 30). Furthermore, significant downregulation in the expression level of *Tnf-α* was also noted, while the expression levels of *Il-6* were moderately suppressed (**Figure 6D**). In contrast, genes associated with anti-inflammatory and tissue repair activities, including *Chil3*, *Arg1*, and *Retnla*, were all noted to be significantly upregulated supporting our findings of more numerous resident cardiac macrophages and M2-polarized monocyte-derived macrophages (**Figure 6E**). Together, our data introduce the therapeutic potential of THP1-IL4-exo, one type of M2-macrophage exosomes, for the treatment of cardiac inflammation and functional decline that develops as a consequence of MI.

## Discussion

Despite decades of research pointing to the involvement of inflammation as a contributor to the pathogenesis of HFrEF post-MI (7, 25), the use of anti-inflammatory drugs targeting NF-kB-signaling have so far failed to show benefits in clinical outcomes (31). Thus, it remains uncertain whether immune modulatory therapies could serve to effectively control HFrEF, which remains a significant source of premature morbidity and death (32). A promising approach could include the selective targeting of the innate immune pathway known as the cGAS-STING-IRF3-T1 IFN axis, a DAMP sensing system that recognizes dsDNA released during ischemic cell death, a phenomenon common to numerous forms of cardiovascular diseases (11, 33). Through a comprehensive series of studies, King et al. revealed that interrupting T1 IFN signaling via antibody-mediated targeting of IFNAR provides a therapeutic avenue to alleviate the onset and severity of HFeEF subsequent to MI (9).

Building on our recent findings that introduced therapeutic benefits of M2-polarized macrophage exosomes in controlling cardiometabolic inflammation, insulin resistance, and atherosclerosis in obese diabetic mice (17, 19), we tested in this study their utility to control adverse cardiac remodeling and HF post-MI. With human translation in mind, we opted to study exosomes produced by the human monocytic cell line THP-1, as we recently reported (17).

Our studies were performed using the ApoE^h/h^/SR-B1^−/−^/Mx1-Cre^+^ mouse model of diet-induced MI and HF that we developed and reported in recent studies (20, 21). The model has since been used by other laboratories and is recognized as one that faithfully mimics human coronary heart disease and MI (34). In this model, expression of hypomorphic ApoE^h/h^ alleles, when paired with a loss of SR-B1 expression, provides a convenient platform to reproducibly generate mice with occlusive coronary atherosclerosis and ischemic myocardial disease within 4 weeks of initiating a diet rich in fat and cholesterol (20, 21).

Subsequent normalization of plasma cholesterol levels, achieved by returning the mice to a chow diet and inducing Cre-mediated repair of the HypoE alleles, enables a deceleration in atherosclerosis while prolonging survival, resulting in mice with ischemic myocardial injury. Mice that survive MI develop cardiac inflammation and soon thereafter impaired left ventricular function that models human HFrEF (20, 21). Our prior studies of the model demonstrated the therapeutic utility of profound immunosuppression using the sphingosine 1 phosphate (S1P) analog FTY720 to improve survival and cardiac function among ApoE^h/h^/SR-B1^−/−^/Mx1-Cre^+^ mice with established ischemic cardiac disease (20, 21). While outcomes of that study did not identify a molecular pathway responsible for the physiological benefits, we now speculate that it could have originated from an attenuation of T1 IFN signaling resulting from S1PR1-mediated suppression of INFAR signaling (35) accounting for our reported observations of reduced levels of *Ccl7* and *Oas1a* mRNA expression in cardiac tissue (21).

In this study, we uncovered that IL4-THP1-exo, a type of exosome produced by M2-polarized human macrophages, can serve as an effective cardioprotective agent in preventing cardiac functional decline and heart failure in response to MI. Through serial echocardiographic measurements of cardiac performance in ApoE^h/h^/SR-B1^−/−^/Mx1-Cre^+^ that experienced diet-induced coronary atherosclerosis and cardiac ischemia, we recorded marked improvements in numerous parameters of cardiac LV function that included FS, EF, SV, and LVESV, which all returned to pre-diet baseline levels as soon as four weeks of triweekly IP administrations of IL4-THP1-exo post MI. Together, our findings demonstrate the potency of IL4-THP1-exo in protecting against the onset of HFrEF and premature death post-MI, while sham-treated mice experienced profound declines in cardiac LV function and longevity.

Clues to appreciate the cardioprotective properties afforded by IL4-THP1-exo stem from our recent observations that reported their ability to suppress hematopoiesis and the number of monocytes and neutrophils in the circulation of mice with hyperlipidemia and diabetes (17, 19). As noted in our prior studies, the control of myelopoiesis originated from within the bone marrow with robust decreases in the numbers of hematopoietic stem cells (HSC) and a suppression in the Granulocyte-Monocyte Progenitor (GMP) lineage that serves as a source of myelopoiesis in response to hyperlipidemia, diabetes, and MI (7, 25). THP1-IL4-exo were also effective in decreasing the numbers of circulating Ly- 6C^lo^ and Ly-6C^hi^ monocytes, and neutrophils. The control of myelopoiesis subsequently led to reduced numbers of inflammatory cells recruited to the heart, supporting the benefits of THP1-IL4-exo in provoking a general control of inflammation in mice that experienced MI.

Beyond limiting the recruitment of inflammatory myeloid cells into the ischemic heart, THP1-IL4-exo likely fostered the differentiation of monocyte-derived macrophages into rcMacs recognized as cells that express the TIM-4 cell-surface antigen (26, 36). Indeed, TIM-4^+^ cardiac macrophages are recognized to exert profound anti-inflammatory properties that resolve cardiac inflammation, including through efferocytosis to clear apoptotic cells and cellular debris (26, 37). Their augmented numbers in cardiac tissue of mice treated with THP1-IL4-exo likely contributed to our observed suppression of cardiac injury and the propagation of anti-inflammatory signals. Furthermore, THP1-IL4-exo fostered the retention of TIM-4^+^/CCR2^-^ rcMacs, a unique population of self-renewing myeloid cells recognized for protective functions that include the suppression of cardiac fibrosis, debris removal, and vascular and lymphatic remodeling in response of MI (26, 38). Together, our findings highlight the capacity for THP1-IL4-exo to effectively limit detrimental myeloid cell activity while promoting anti-inflammatory properties following ischemic myocardial injury caused by occlusive coronary atherosclerosis.

Beyond suppressing populations of inflammatory cells in the BM, the circulation, and the heart, THP1- IL4-exo profoundly reprogrammed circulating Ly-6C^hi^ monocytes towards an M2 anti-inflammatory phenotype. Through an unbiased approached designed to identify mechanisms responsible for the therapeutic benefits exerted by THP1-IL4-exo, we identified T1 IFN signaling as the dominant immune pathway controlled in myeloid cells of the circulation, as well as in the bone marrow and cardiac tissue. However, beyond emulating results obtained using antibody-mediated attenuation of INFAR (9), a constellation of yet to be identified cell-signaling properties inherent to THP1-IL4-exo likely contributed to restore cardiac function in mice that experienced MI. Other impacted pathways that could have contributed to improve cardiac healing could include those that enhance oxidative phosphorylation, which is recognized to drive M2 macrophage biology (17, 19). Modulation of genes coding for redox stress control, vascular repair, and suppression of genes known for adverse ventricular modeling, such as MMPs (29, 30), could all serve to alleviate cardiac injury post-MI. These findings hint at the capacity of THP1-IL4-exo to impact a variety of distinct pathways that together foster the restoration of cardiac performance following ischemic myocardial injury.

While a complete understanding of the therapeutic benefits exerted by THP1-IL4-exo will require further study, it is likely that they derive in part from their ability to communicate microRNA to cardiac tissue, similar to what has been reported for microRNA-146a enriched in cardiosphere-cell-derived exosomes (39) and microRNA-21 enriched exosomes produced by cultured mesenchymal stem cells (40, 41). Interestingly, our prior findings documented the presence of these and other anti-inflammatory and metabolic microRNA in THP1-IL4-exo (17). As such, it is likely that in this study, THP1-IL4-exo robustly polarized macrophages in treated mice into an M2 phenotype that inhibited ISG gene expression and fostered the retention and accumulation of TIM-4^+^ rcMacs in cardiac tissue.

Future studies will be required to investigate whether treatments with THP1-IL4-exo also benefit other cell types within the heart, including cardiomyocytes and fibroblasts, to improve cardiac function.

Furthermore, based on our observations demonstrating a capacity for THP1-IL4-exo to improve mitochondrial metabolism while controlling oxidative stress (17), similar benefits could have led to improved cardiomyocyte mitochondrial health and viability. Additional studies will also be required to investigate the ability for THP1-IL4-exo to communicate benefits to atheroma in coronary arteries to drive the remodeling and stabilization of atherosclerotic plaques. Finally, it will be both interesting and of value to test whether THP1-IL4-exo can provide therapeutic benefits for other forms of HF, including those that develop as a consequence of pulmonary hypertension, diabetic heart failure with preserved ejection fraction (HFPEF), and aging.

## Acknowledgements

We thank the staff of the Electron Microscope Laboratory at the University of California, Berkeley for advice and assistance with electron microscopy sample preparation and imaging.

## Author contributions

Conceptualization, M.N. and R.L.R.; methodology, M.N., A.S.G., and R.L.R.; investigation, M.N., A.S.G., T.A.P., N.K.V., and R.L.R.; visualization, M.N., A.S.G., and T.A.P.; funding acquisition, R.L.R; project administration, R.L.R.; supervision, R.L.R.; writing – original draft, M.N. and R.L.R.; writing – review & editing, M.N., A.S.G., and T.A.P., N.K.V., and R.L.R.

